# Effects of early life glucocorticoid exposure on metabolism in zebrafish (*Danio rerio*) larvae

**DOI:** 10.1101/2019.12.16.877654

**Authors:** Ruud van den Bos

## Abstract

In this study we assessed the effects of increased cortisol levels during early embryonic development (0-6 hours post-fertilisation (hpf)), thereby mimicking maternal stress, on metabolism in zebrafish (*Danio rerio*) larvae. In two series of experiments fertilized eggs were exposed to a cortisol-containing, a dexamethasone-containing (to stimulate the glucocorticoid receptor (GR) specifically) or a control medium for 6 hours post-fertilisation (0-6 hpf). In the first series we measured oxygen consumption as a proxy for metabolism, in the second series gene-expression of genes related to gluconeogenesis and glucose transport. Previously we have found that at 5 days post-fertilisation (dpf) baseline cortisol levels are increased following cortisol pre-treatment but not following dexamethasone pre-treatment, suggesting a higher hypothalamus-pituitary-interrenal cells (HPI-axis) activity. Hence, we hypothesized that oxygen consumption and gene-expression were stronger in cortisol-treated than in dexamethasone-treated and control-treated subjects at 5 dpf. Indeed, we observed increased oxygen consumption in cortisol-treated subjects compared to dexamethasone-treated or control-treated subjects. However, gene-expression levels were not different between treatments, which may have been due to a developmental delay in this second series. We also reasoned that both cortisol-treated and dexamethasone-treated subjects would show a higher metabolism at 1 dpf than control-treated subjects as the HPI-axis is not functional as yet and more general processes are being stimulated by cortisol through GR stimulation. Indeed, we observed increased oxygen consumption and increased expression of genes related to gluconeogenesis and glucose transport in cortisol-treated and dexamethasone-treated subjects than control-treated subjects. These data show that early-life exposure to cortisol, mimicking thereby maternal stress, increased metabolism at different life stages, i.e. 1 and 5 dpf, involving the GR.

## 1. Introduction

In zebrafish, mothers deposit, among others, cortisol and mRNA of the glucocorticoid receptor (GR), in oocytes; cortisol levels decrease across 24 hours after which zygotes start to produce cortisol by the then developing interrenal cells (Liu, 2007; Pikulkaew *et al.*, 2010, 2011; Wilson *et al.*, 2013). It does not take until after hatching (48-72 hrs) before pituitary control over interrenal cortisol production occurs, after which it takes another 4-5 days before the axis is fully functionally mature (reviews: Alsop and Vijayan, 2008; Alderman and Bernier, 2009). Maternal GR mRNA is present during the first 6 hrs; from 8-9 hrs onwards zygotic expression of GR commences, while the mineralocorticoid receptor (MR) is expressed from 24 hrs onwards (Alsop and Vijayan, 2008; Pikulkaew *et al.*, 2010, 2011).

As a means of mimicking maternal stress, levels of cortisol have been enhanced by injections of cortisol into the yolk of one-cell stage embryos (e.g. Best et al., 2017; Nesan and Vijayan, 2012, 2016) or through exposure by the medium (Hartig et al., 2016; van den Bos *et al*., 2019a, 2019b). These studies have shown that baseline-levels of cortisol are increased following treatment in larvae (Best *et al.*, 2017; Nesan and Vijayan, 2012; van den Bos *et al.*, 2019a). Here, we measure whether this in turn increases metabolism measuring oxygen consumption and gene-expression of genes involved in gluconeogenesis and glucose transport.

In earlier experiments we have found that at 5 days post-fertilisation (dpf) baseline cortisol levels are increased following cortisol treatment for 6 hours post fertilisation (hpf) but not following dexamethasone (a potent GR-agonist (Rupprecht *et al.*, 1993) to assess the role of GR more specifically) treatment (Althuizen, 2018; van den Bos *et al.*, 2019a), suggesting a higher hypothalamus-pituitary-interrenal cells (HPI-axis) activity. Higher cortisol levels have been shown to be associated with increased respiration rate (oxygen consumption and CO_2_ production) and plasma glucose levels through stimulation of GR (De Boeck et al., 2001; Jimeno et al., 2017, 2018; Wendelaar Bonga, 1997). Hence, we hypothesized that oxygen consumption and gene-expression were stronger in cortisol-treated than in control-treated or dexamethasone-treated subjects at 5 dpf.

Oxygen consumption was measured using a non-invasive automated optical sensor system. For glucose, we assessed mRNA expression levels of genes involved in gluconeogenesis and glucose transport: *glucose-6-phosphatase (g6pca), solute carrier family 5 member 1 (slc5a1), phosphoenolpyruvate carboxykinase 1 (pck1)* and *phosphoenolpyruvate carboxykinase 2 (pck2). Pck1* and *pck2* are involved in the first (rate-limiting) steps of gluconeogenesis, *slc5a1* is a membrane transporter protein involved in the active transport of glucose into cells, while *g6pca* is involved in the final step of gluconeogenesis (Chatzopoulou *et al.*, 2015; Willi *et al.*, 2018; Zhao *et al.*, 2016). *Pck1* is more dynamically involved in gluconeogenesis and glucose regulation than *pck2* and hence differences may be more prominently found in *pck1* than *pck2* (Jurczyk *et al.*, 2011). Previous research has shown that exposure to glucocorticoids increased the transcript abundance of these genes in zebrafish larvae (Chatzopoulou *et al.*, 2015; Elo *et al.*, 2007; Hartig *et al.*, 2016; Willi *et al.*, 2018, 2019; Zhao *et al.*, 2016).

As zygotic cortisol production develops from 24 hours onwards (Liu, 2007; Pikulkaew *et al.*, 2010, 2011; Wilson *et al.*, 2013) and hence differences in endogenous base-line production of cortisol may be small at this time-point, we reasoned that at 24 hours (1 dpf) cortisol-treatment and dexamethasone treatment would produce similar effects: both are stimulating GR in early life increasing thereby metabolism.

## 2. Materials and Methods

### 2.1. Subjects, spawning and care

Housing conditions and breeding procedures were similar as those reported in van den Bos *et al.* (2019a, 2019b). In short, in-house bred adult (> 6 months) zebrafish of the AB strain from the fish facilities of the Department of Animal Ecology and Physiology (Radboud University, Nijmegen, the Netherlands) were used for egg production. They were kept in recirculation systems (bio-filtered Nijmegen tap water, ~28°C, pH 7.5-8, conductivity ~320 microSiemens/cm; Fleuren and Nooijen, Nederweert, the Netherlands) in 2-litre aquaria (approximately 30 fish of mixed sex) under a 14h:10h light-dark cycle (lights on from 09.00h to 23.00h) fed twice daily (at 09.00 h (*Artemia* sp. and Gemma Micro 300 (Skretting, Wincham, Northwich, Cheshire, UK)) and 15.00 h (Gemma Micro 300 (Skretting, Wincham, Northwich, Cheshire, UK)).

Breeding started at least one hour after the last feeding of zebrafish (>16.00 h). Two males and one female of the AB strain were placed in a zebrafish breeding tank, separated by a partitioning wall, with water of ~28°C and an artificial plant on the side of the female. Fish were placed in the dark until the next morning. After turning on the lights at 09.00h, fish were allowed to acclimatise for a few min. Then, water was changed for clean warm water of ~28°C, the partitioning wall was removed and tanks were placed at a slight angle, such that the fish had the possibility to move into shallow water to spawn. When no eggs were produced, water was once more renewed and tanks were again placed at a slight angle.

### 2.2. Exposure

Cortisol (hydrocortisone; Sigma-Aldrich, Zwijndrecht, the Netherlands) and dexamethasone (Sigma-Aldrich, Zwijndrecht, the Netherlands) were dissolved in 96% ethanol in the required stock solution concentrations and stored at −20°C. For each experiment solutions with the appropriate concentration were freshly prepared from these stock solutions (Althuizen, 2018; van den Bos *et al.*, 2019a, 2019b): cortisol-containing medium: 400 μg/l cortisol (1.1 μM), 0.4 ml/l 96% ethanol, 5 mM NaCl, 0.17 mM KCl, 0.33 mM CaCl_2_, 0.33 mM MgSO_4_ in dH2O; dexamethasone-containing medium: 430 μg/l dexamethasone (1.1 μM), 0.4 ml/l 96% ethanol, 5 mM NaCl, 0.17 mM KCl, 0.33 mM CaCl_2_, 0.33 mM MgSO_4_ in dH2O. Control medium consisted of: 0.4 ml/l 96% ethanol, 5 mM NaCl, 0.17 mM KCl, 0.33 mM CaCl_2_, 0.33 mM MgSO_4_ in dH2O.

Directly following spawning and fertilization eggs were collected and exposed to the solutions. Eggs of each spawning were randomly assigned to a Petri dish filled with 25 ml of either control medium, cortisol-containing medium or dexamethasone-containing medium.

Within 1-1.5 hour post fertilisation (hpf) Petri dishes were placed in an incubator set at 28.5 °C (300–350 lux). Eggs were exposed to these solutions for 6 hrs (they take up water from the environment in this period and swell). It has been shown that both cortisol and dexamethasone diffuse inside the eggs in this period (Steenbergen *et al.*, 2017). In addition we have shown this procedure to be effective in eliciting changes in physiology, behaviour and immune function at 3-5 days post fertilisation (dpf; Althuizen, 2018; van den Bos *et al.*, 2019a, 2019b).

Following this, cortisol-containing, dexamethasone-containing and control media were replaced by E3 medium (5 mM NaCl, 0.17 mM KCl, 0.33 mM CaCl_2_, 0.33 mM MgSO_4_, 3 ml/l 0.01% (w/v) methylene blue, in dH2O) three times to ensure that the original treatment media were completely removed. Petri dishes were returned to the incubator for the embryos to develop further (28.5°C; 14h:10h light-dark period (lights on: 09.00h – 23.00h); light phase: 300–350 lux; dark phase; 0 lux). At 1 dpf and 4 dpf E3 medium was refreshed and unfertilized eggs, dead eggs/embryos/larvae and chorions were removed.

Two series were run: one series was used for O_2_ measurements (and immune-related experiments reported elsewhere; van den Bos *et al.*, 2019b) and one series was used for gene-expression (and immune-related experiments reported elsewhere; van den Bos *et al.*, 2019b).

All experiments were carried out in accordance with the Dutch Experiments on Animals Act (http://wetten.overheid.nl/BWBR0003081/2014-12-18), the European guidelines for animal experiments (Directive 2010/63/EU; http://eur-lex.europa.eu/legal-content/NL/TXT/HTML/?uri=CELEX:32010L0063) and institutional regulations. Larvae were euthanized by placing them in ice slurry for at least 20 minutes followed by adding bleach to the slurry.

### 2.3 Oxygen consumption

Oxygen consumption of larvae exposed to either cortisol-containing medium, dexamethasone-containing medium or control medium was measured with the SDR Sensordish® Reader version 4.0.0 (PreSens, Regensburg, Germany).

A 24-well SensorDish (PreSens, Regensburg, Germany) was submerged in a tank containing E2 and any air bubbles, which could interfere with reliable measurements, were forced out of the wells by pipetting. Seven embryos (1 dpf) or larvae (5 dpf) of each treatment (cortisol, dexamethasone, control) were added to the wells (one embryo/larva per well; 200μl/well) and three wells were filled with E2 as blanks. After loading wells were sealed with Microseal® ‘B’ seal (Bio-Rad Laboratories, the United Kingdom) avoiding that air bubbles, which could interfere with the measurements, were present; if this happened, this well was discarded from analysis.

The 24-well plate was placed in the SDR SensorDish Reader and recordings were done running the software programme MicroResp® version 1.0.4 (Loligo Systems, Viborg, Denmark). The Grant LT Ecocool 150® (Grant Instruments, Cambridge, United Kingdom) created a continuous flow of water (27°C) over the SensorDish. Values were adjusted for the atmospheric pressure of the specific recording day (atmospheric pressure of Nijmegen obtained through the internet). Recordings started at 11:00 AM; oxygen was measured every 30 seconds for 5 hours.

### 2.3 Gene-expression using qPCR

Cortisol-treated, dexamethasone-treated and control embryos or larvae were sampled from the Petri dishes at 3:00 PM for analysis of gene expression. Four embryos or three larvae were transferred to a 2ml Eppendorf tube and the residual medium was removed with a pipette; so each sample contained material of more subjects. Tubes were subsequently snap frozen in liquid nitrogen, kept on ice during further sampling and afterwards stored at −80°C until total RNA extraction. RNA isolation, RNA preparation, removal of genomic DNA from the samples and synthesis of the cDNA were performed according to the protocol described in detail in van den Bos *et al.* (2019a, 2019b). Analysis of the data was carried out using a normalisation index of two reference genes (viz. *elongation factor alpha* (*elf1a*) and *ribosomal protein L13* (*rpl13*)) (Vandesompele *et al.*, 2002). Primer sequences of genes of interest are listed in Table 1. Next to genes involved in gluconeogenesis, a few selected HPI-axis genes were included as control for the effectiveness of the procedure (Althuizen, 2018; van den Bos *et al.*, 2019a).

**Table 1.**
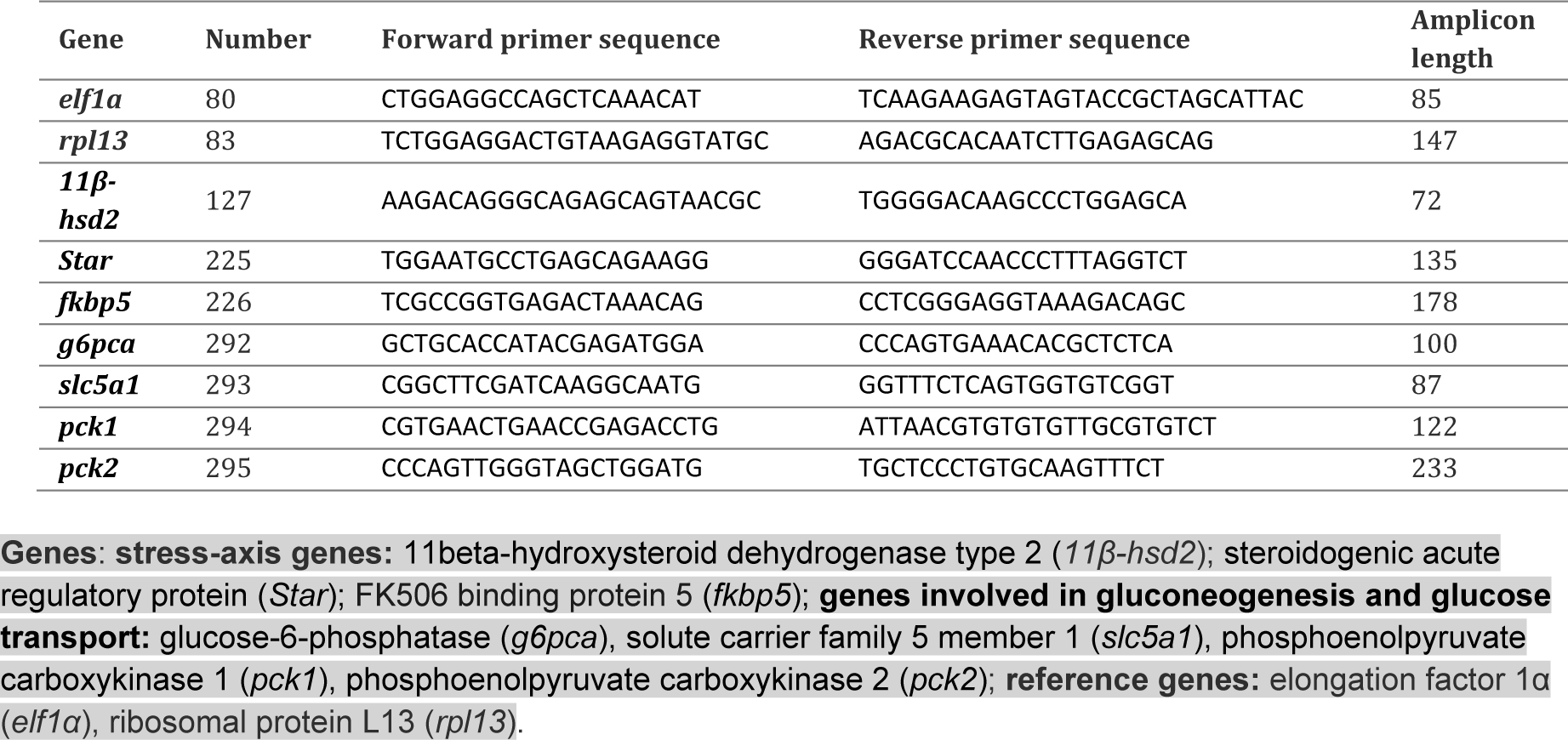
Nucleotide sequences of the forward and reverse primers used for qPCR.

### 2.5 Data analysis and statistics

For oxygen consumption data were first corrected by the values measured in the empty wells (blanks). The means of the n=3 blanks were calculated per 30 seconds. At each time-point the measurements of the experimental wells were corrected by the means of the blanks according to the formula: (value well/mean value blanks) *100. Values are expressed as percentage oxygen present in the well. To limit the number of time points for analysis only values at 10-min intervals were considered. Oxygen data were analysed using a two-way analysis of variance (ANOVA; with treatment as factor and time as repeated measurement) followed by post-hoc Tukey HSD where appropriate.

For gene expression outliers were removed following a Grubb’s test (p≤0.01). Differences between treatments were analysed using a one-way ANOVA (treatment as factor) followed by post-hoc Tukey HSD. All tests were done using IBM SPSS version 23 (IBM, Armonk, NY, USA). Significance was accepted when p≤0.05 (two-tailed) and trends are indicated (p≤0.10) where appropriate; ns: p>0.10.

## 3. Results

### 3.1. Expression of HPI-axis genes at 1 dpf and 5 dpf

Table 2 shows transcript abundance of a number of selected genes of the HPI-axis. At 1 dpf, but not 5 dpf, the expression of *Star* was lower in the cortisol-treated group compared to the control-treated group. At both 1 dpf and 5 dpf transcript abundance of *fkbp5* was significantly increased following dexamethasone treatment compared to control treatment. At 1 dpf, but not 5 dpf, transcript abundance of *11β-HSD2* was significantly increased following dexamethasone treatment compared to control treatment.

**Table 2:** Mean (±SEM) transcript abundance of selected genes involved in the HPI-axis at 1 dpf and 5 dpf. Capitals indicate which groups are similar (post-hoc Tukey HSD; p≤0.05) following a significant one-way ANOVA. For *11β-HSD2* at 5 dpf a trend was found (F(2,20)=3.022, p≤0.071).

|  | 1 dpf |  |  |
| --- | --- | --- | --- |
| gene | control (n=8) | cortisol (n=8) | dexamethasone (n=8) |
| *Star* | 0.093±0.007<sup>A</sup> | 0.059±0.010<sup>B</sup> | 0.086±0.008<sup>A,B</sup> |
| *11β-HSD2* | 0.069±0.003<sup>A</sup> | 0.054±0.003<sup>A</sup> | 0.096±0.012<sup>B</sup> |
| *fkbp5* | 0.0078±0.0003<sup>A</sup> | 0.0080±0.0005<sup>A</sup> | 0.0573±0.0032<sup>B</sup> |
|  | 5 dpf |  |  |
| gene | control (n=8) | cortisol (n=7) | dexamethasone (n=8) |
| *Star* | 0.95±0.06 | 0.87±0.05 | 0.93±0.04 |
| *11β-HSD2* | 0.74±0.06 | 0.70±0.06 | 0.89±0.05 |
| *fkbp5* | 0.31±0.06<sup>A</sup> | 0.37±0.05<sup>A</sup> | 0.74±0.09<sup>B</sup> |

### 3.2. Metabolism of embryos at 1 dpf

Oxygen consumption at 1 dpf after the different 0-6 hpf treatments is shown in figure 1: the amount of oxygen decreased with 35% across a 5-hour time span and groups exposed to either cortisol or dexamethasone had lower %oxygen-values than the control group; an effect which became stronger as time progressed. A repeated measures ANOVA yielded a significant difference across time (F(30,540)=601.580, p<0.001), a significant interaction term (F(60,540) =2.457, p<0.001) and a significant effect of treatment (F(2,18)=3.937, p≤0.038). Both the cortisol-treated group (Tukey HSD: p≤0.049) and the dexamethasone-treated group (Tukey HSD: p≤0.085) had overall lower %oxygen-values than the control-treated group. This suggests that oxygen consumption was higher in glucocorticoid-treated subjects than in control-treated subjects.

**Figure 1:**
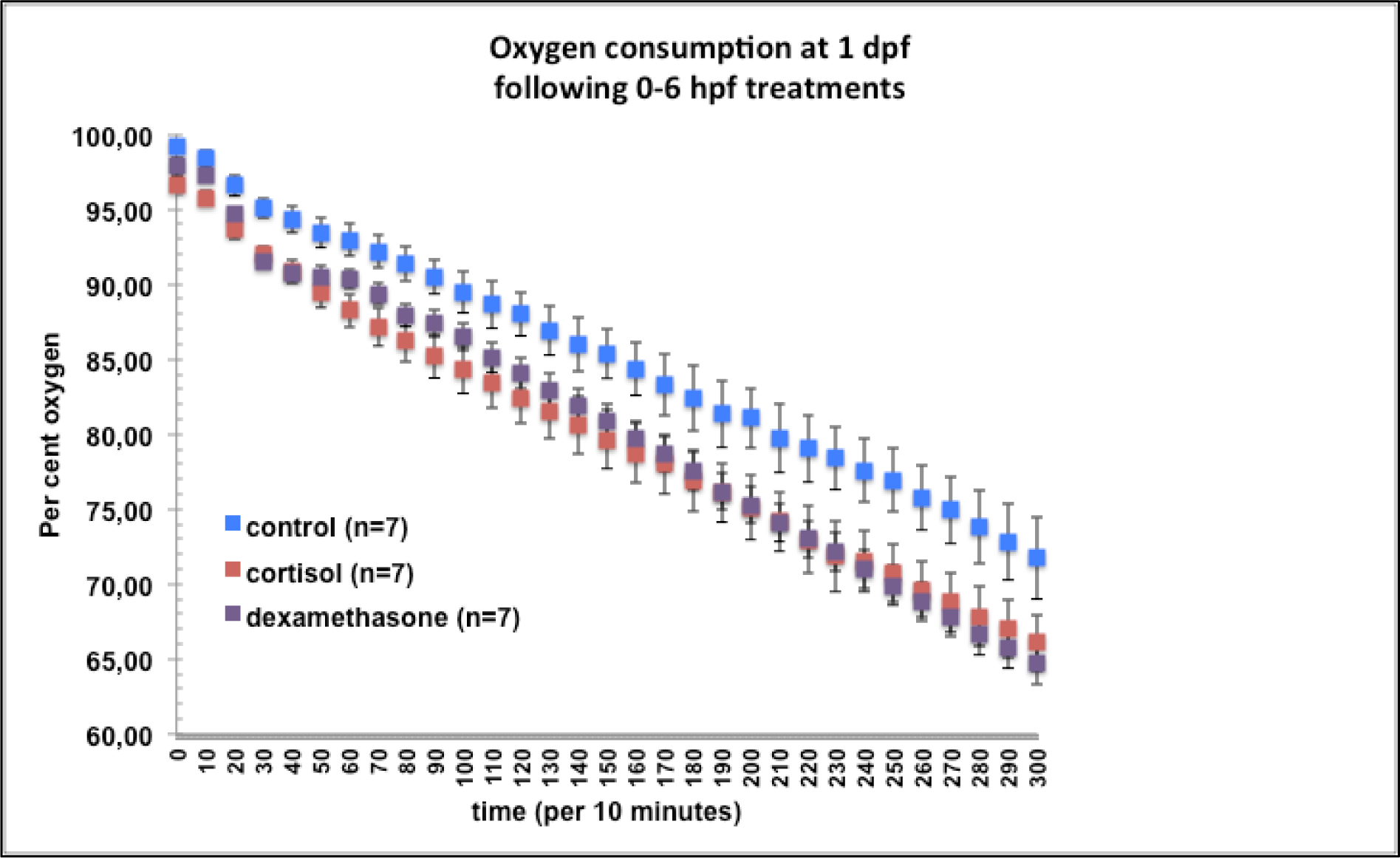
Mean (± SEM) oxygen values (%) at 1 dpf for the different 0-6 hpf treatments. Note that the y-axis starts at 60% oxygen.

Figure 2 (panels A and B) shows the transcript abundance of genes involved in gluconeogenesis and glucose transport. Transcript abundance of *pck1*, *g6pca* and *slc5a1* increased following glucocorticoid treatment at 0-6 hpf compared to control-treated subjects, particularly in cortisol-treated subjects.

**Figure 2:**
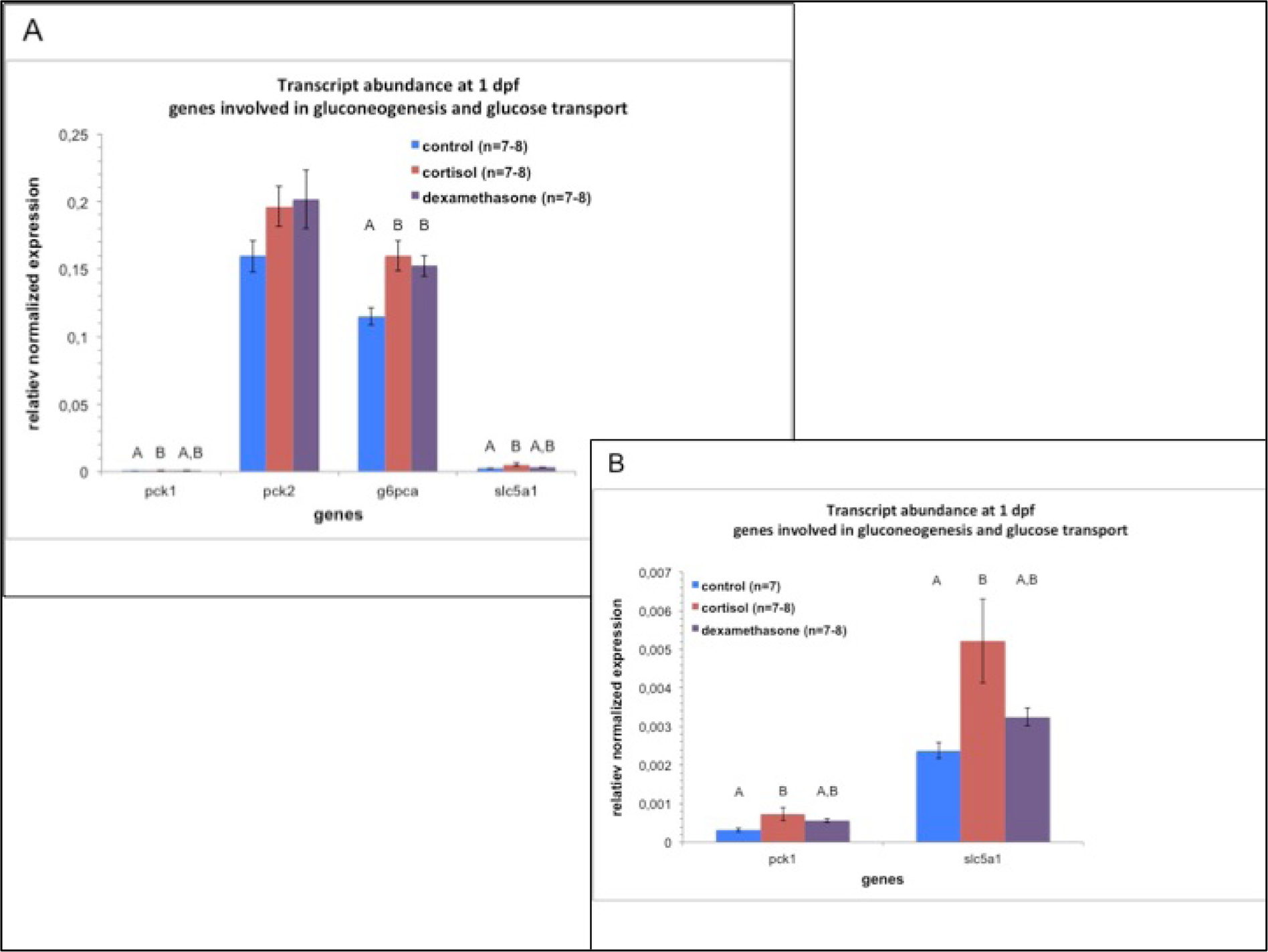
Mean (±SEM) transcript abundance of genes involved in gluconeogenesis and glucose transport at 1 dpf. Capitals indicate which groups are similar (post-hoc Tukey HSD; p≤0.05) following a significant one-way ANOVA. Panel A shows transcript abundance of all genes, panel B of *pck1* and *slc5a1*, whose values were low and therefore difficult to see in panel A. The following values were eliminated after Grubb’s test (p≤0.01): control: *pck1* (0.00219) and *slc5a1* (0.0063); dexamethasone: *slc5a1* (0.0245). For cortisol *pck1*: n=7 samples.

### 3.3 Metabolism of larvae at 5 dpf

Oxygen consumption at 5 dpf after the different 0-6 hpf treatments is shown in figure 3: the amount of oxygen decreased over time and after five hours almost 80% of the available oxygen was consumed. In particular the curve of the cortisol-treated subjects started to level off towards the end of the recordings, reaching floor values. The cortisol-treated group had lower %oxygen-values than the dexamethasone-treated group or the control-treated group; an effect that became stronger as time progressed. A repeated measures ANOVA indicated a significant difference across time (F(30,540)=2017.862, p<0.001), a significant interaction term (F(60,540)=3.416, p<0.001) and a significant effect of treatment (F(2,18)=5.786, p≤0.011). The cortisol-treated group had overall lower %oxygen-values that the dexamethasone-treated group (Tukey HSD p≤0.009), while no overall significant difference was found with the control-treated group. As the effect of cortisol treatment was most prominent towards the end of the recordings, the last 2 hours (from 190 until 300 minutes) were analysed separately. A repeated measures ANOVA indicated a significant difference across time (F(11,198)=300.525, p<0.001), a significant interaction term (F(22,198)=3.393, p<0.001) and a significant effect of treatment (F(2,18)=6.917, p<0.006). The cortisol-treated group had overall lower %oxygen-values than both the control-treated group (Tukey HSD: p≤0.085) and the dexamethasone-treated group (Tukey HSD: p≤0.005) during this time-window. This suggests that oxygen consumption was higher in cortisol-treated subjects than in control-treated or dexamethasone-treated subjects.

**Figure 3:**
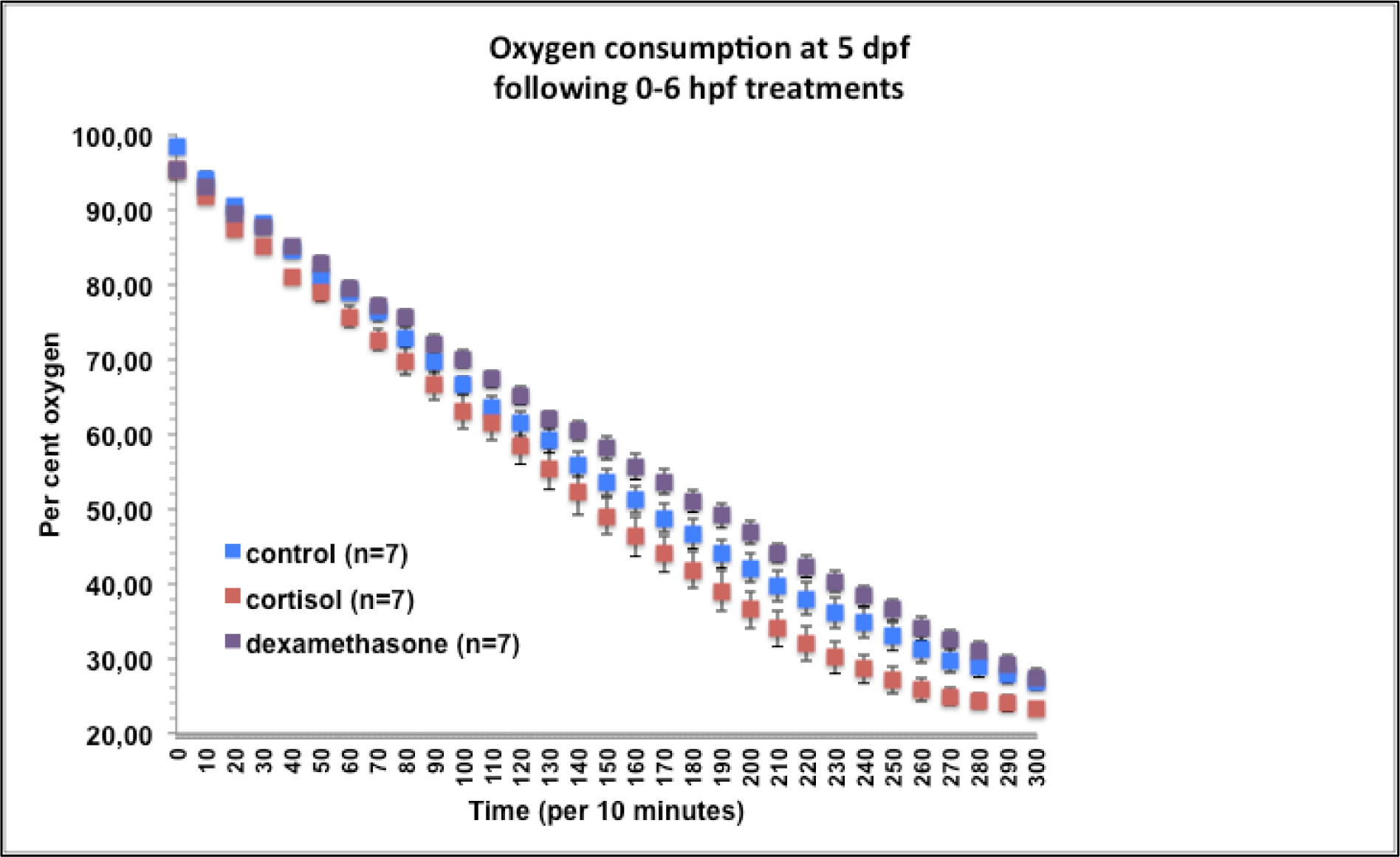
Mean (± SEM) oxygen values (%) at 5 dpf for the different 0-6 hpf treatments. Note that the y-axis starts at 20% oxygen

Figure 4 (panels A and B) shows the transcript abundance of genes involved in gluconeogenesis and glucose transport. No effects were found at 5 dpf. However it should be noted that transcript abundance of *pck*1 was low at 5 dpf (see Discussion).

**Figure 4:**
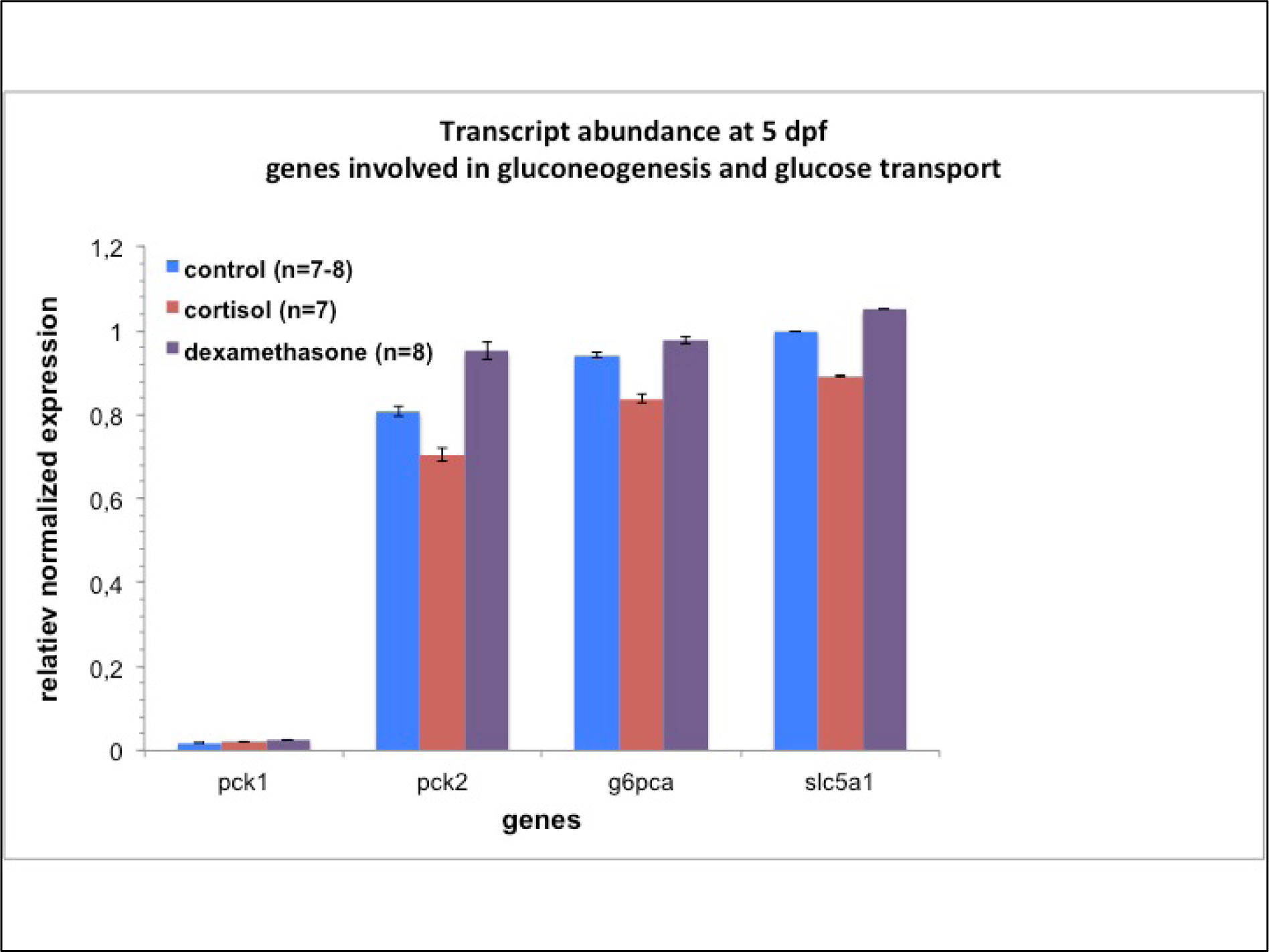
Mean (±SEM) transcript abundance of genes involved in gluconeogenesis and glucose transport at 5 dpf. For cortisol n=7 samples; for control: *pck1:* n=7 samples.

## 4. Discussion

The data of this study showed that cortisol treatment at 0-6 hpf, mimicking thereby maternal stress, increased metabolism at both 1 dpf and 5 dpf, and that GR is involved.

### General

To assess whether our procedure of exposing embryos to glucocorticoid medium for 6 hours was effective we measured transcript abundance of a number of selected genes as markers (Althuizen, 2018; van den Bos *et al.*, 2019a). In line with earlier findings dexamethasone, but not cortisol, strongly increased transcript abundance of FK506 binding protein 5 (*fkbp5*) both at 1 dpf and 5 dpf (Althuizen, 2018; van den Bos *et al.*, 2019a). Fkbp5 is a GR responsive gene that is up-regulated under high levels of cortisol, thereby down-regulating GR activity (Zannas *et al.*, 2016). Thus *fkbp5* expression is a marker for levels of GR stimulation (e.g. Willi *et al.*, 2018). As we used equimolar doses of cortisol and dexamethasone (1.1 μM) and dexamethasone is more potent than cortisol at GR (Rupprecht *et al.*, 1993), we may have stimulated GR more strongly following dexamethasone than cortisol exposure.

In earlier studies we observed increased transcript abundance of *11β-HSD2* after dexamethasone exposure (Althuizen, 2018), but not cortisol exposure (van den Bos *et al.*, 2019a), which was confirmed here at 1 dpf, but only partially at 5 dpf. 11β-HSD2 inactivates cortisol by metabolizing cortisol into the inactive metabolite cortisone (Alsop and Vijayan, 2008; Alderman and Bernier, 2009; Wendelaar Bonga, 1997) and its activity is enhanced after GR stimulation to control cortisol levels (e.g. Draper and Stewart, 2005; Wilson *et al.*, 2016). As indicated above dexamethasone treatment activated GR more strongly than cortisol treatment and hence we observed an effect on the expression of *11β-HSD2*. This finding may underlie why we did not observe increased cortisol levels at 5 dpf following dexamethasone exposure (Althuizen, 2018). Even before the HPI-axis starts to function fully (Alsop and Vijayan, 2008; Alderman and Bernier, 2009) the enzyme that inactivates cortisol is already expressed at higher levels. Earlier we also observed at 5 dpf that dexamethasone (Althuizen, 2018), but not cortisol (van den Bos *et al.*, 2019a), decreased transcript abundance of *11β-hydroxylase*, the gene coding for the final and crucial enzyme in cortisol synthesis (Alsop and Vijayan, 2008; Alderman and Bernier, 2009; Wendelaar Bonga, 1997), also supportive of the observation that we did not observe an increase of cortisol levels following dexamethasone exposure.

In line with earlier studies (Althuizen, 2018; van den Bos *et al.*, 2019a) we did not observe changes in transcript abundance of *Star* at 5 dpf. However, we observed a decrease at 1 dpf following cortisol but not dexamethasone treatment. Steroidogenic acute regulatory protein regulates cholesterol shuttling across the mitochondrial membrane, which is one of the rate-limiting steps in steroid synthesis (Liu, 2007). Steroidogenesis commences at 28 h after fertilization, when, among other genes, *Star* is expressed (Liu, 2007; Nesan and Vijayan, 2013). Maternal cortisol levels decrease across 24 hours after which zygotes start to produce cortisol by the then developing interrenal cells (Liu, 2007; Pikulkaew *et al.*, 2010, 2011; Wilson *et al.*, 2013). Whether higher levels of maternal cortisol (as induced here by exposure to cortisol) slightly delay the onset of *Star* expression and hence the onset of zygotic cortisol production awaits further studies.

Oxygen consumption was overall higher in 5 dpf larvae than in 1 dpf embryos in line with earlier studies (Huang *et al.*, 2013; Makky *et al.*, 2008). We noted that *pck1* expression was rather low in the 5 dpf larvae. At this stage *pck1* is primarily found in the liver to dynamically regulate glucose levels (Jurczyk *et al.*, 2011). As yolk nutrients are being consumed, caloric deficits increase with up-regulation of *pck1 (*‘feeding to fasting transition’: Gut *et al.*, 2013; Rocha et al., 2014). Hence, the relatively low levels that we observed may indicate that these larvae still rely on the yolk sac for nutrition. We also measured gene-expression of a few samples (n=3) of the first series (the series in which we measured oxygen consumption) and the expression level of this series was substantially higher, suggesting more advanced development or depletion of the yolk sac (0.26±0.25 (n=3) versus 0.018±0.008 (n=7)).

### Metabolism of embryos at 1 dpf

In line with our hypothesis, oxygen consumption at 1 dpf was enhanced following 0-6 hpf glucocorticoid treatment suggesting a higher metabolic rate as a consequence of stimulating GR. In line with this gene-expression of genes involved in gluconeogenesis and glucose transport was enhanced (see also Chatzopoulou *et al.*, 2015; Elo *et al.*, 2007; Hartig *et al.*, 2016; Willi et al., 2018, 2019a; Zhao *et al.*, 2016). In the early larval stages *pck1* has been shown to be expressed in the yolk syncytial layer, playing a role in transfer of nutrients from yolk to embryo (conversion of amino acids to glucose) and actively developing tissue like the tail bud, eye and (mid)brain, i.e. it provides local glucose where it is needed (Jurczyk *et al.*, 2011). Glucose has been shown to peak at 24 hpf (Rocha et al., 2014; Jurczyk *et al.*, 2011). Thus the increased expression of *pck1* following glucocorticoid pre-treatment, especially following cortisol, may enhance this peak of glucose and lead to developmental changes.

The functional consequences of enhanced gluconeogenesis are not clear as yet and therefore only speculative. For instance, it has been shown that increasing levels of glucose through the medium is associated with enhanced levels of cortisol in zebrafish (Powers *et al.*, 2010), which may occur through local interrenal glucose sensing mechanisms (Conde-Sieira *et al.*, 2013; for discussion see: Flik and Gorissen, 2016). To what extent the increased baseline levels of cortisol following cortisol-treatment (van den Bos *et al.*, 2019a) are related to this remains to be studied.

### Metabolism of larvae at 5 dpf

In line with our hypothesis, oxygen consumption at 5 dpf was enhanced following 0-6 hpf cortisol but not dexamethasone treatment suggesting a higher metabolic rate. It should be noted that we have observed in earlier experiments that our treatments did not lead to changes (cortisol: van den Bos *et al.*, 2019a) or only slight decreases (dexamethasone (2%); Althuizen, 2018) in body length of larvae. So, while the data of cortisol-treated larvae may not be explained by changes in body size, the slightly lower consumption of dexamethasone-treated subjects potentially could. Regarding dexamethasone, an alternative explanation may be that larvae have lower levels of cortisol. Indeed, we have noted slightly lower levels of cortisol than controls at 5 dpf following 0-6 hpf dexamethasone treatment (Althuizen, 2018).

Earlier we have observed that following cortisol (van den Bos *et al.*, 2019a), but not dexamethasone (Althuizen, 2019a) treatment, baseline levels of cortisol were higher, suggesting higher baseline HPI-axis activity. Higher levels of cortisol have been shown to be associated with higher levels of oxygen consumption or metabolic rate (De Boeck *et al.*, 2001; Jimeno *et al.*, 2017, 2018), which could explain the data. As this is normally also associated with increased levels of glucose through stimulation of GR (De Boeck *et al.*, 2001; Jimeno *et al.*, 2017, 2018; Wendelaar Bonga 1997) we expected increased expression of genes related to gluconeogenesis and glucose transport. However, we did not observe such an effect.

As indicated above it seemed that the series where we obtained the samples for gene-expression was lagging behind in development compared to the series where we measured oxygen consumption. It may be reasoned that the ‘feeding to fasting transition’ when yolk nutrients become depleted (Gut *et al.*, 2013) may strengthen differences between groups. We measured a limited number of samples of the first series for gene expression of *pck1* following cortisol or dexamethasone treatment and observed a stronger increase in cortisol-treated subjects than in control-treated or dexamethasone-treated subjects: cortisol: 0.87±0.28 (n=3), dexamethasone: 0.56±0.21 (n=4), control: 0.26±0.25 (n=3). It is clear however that more studies are needed to substantiate this.

### Limitations

A clear limitation is that we only used one dose of cortisol and dexamethasone. For convenience we used equimolar doses of cortisol and dexamethasone, which may have led to different levels of stimulation of GR 0-6 hpf and consequently (slightly) different effects. Thus, in future studies different dose-ranges may be warranted.

While we measured gene-expression of genes involved in gluconeogenesis and glucose transport we did not measure glucose or other relevant compounds (Chatzopoulou *et al.*, 2015). Future studies should involve these parameters as well to obtain a more complete picture, although gene-expression of the gluconeogenesis-related or transport-related genes seems a valid proxy for glucose levels (Chatzopoulou *et al.*, 2015).

Finally, and not unprecedented (van den Bos *et al.*, 2019b), we noted differences between experimental series. As discussed elsewhere (van den Bos *et al.*, 2019c) this is a general issue while using zebrafish in research as many factors affect early-life development ranging from genetic factors to environmental factors, which are not always fully controllable. Hence, only through repeated experiments and using different read-outs a coherent picture emerges that generates new testable hypotheses. The experiments as described here are another step in this process.

### Conclusion

In conclusion, the present data show that early-life exposure to cortisol, mimicking thereby maternal stress, induces increased metabolism in larvae, which may be related to an increased HPI-axis activity. While this may be adaptive on the short term to fuel processes enhancing survival, it remains to be studied to what extent this may lead to metabolic disorders, such as obesity, hyperglycaemia or type-2 diabetes as a long term consequence of high levels of glucose (Chatzopoulou *et al.*, 2015; Wilson *et al.*, 2016).

## 5. Acknowledgements

I would like to acknowledge the contribution of the following people: Benjamin Eggers and Wies van Zwieten for carrying out experiments as part of their BSc projects; Jan Zethof for carrying out qPCR analysis; Iris van de Pol for assistance in the early phases of the oxygen measurements and Nedim Tüzün (KU Leuven, Belgium) for allowing us to use the oxygen measurement device; Tom Spanings, Antoon van der Horst and Jeroen Boerrigter for excellent fish care.

